# Glucose-dependent metabolism of hippocampal primary neurons in response to chemically induced long-term potentiation

**DOI:** 10.1101/2025.02.24.639988

**Authors:** Natalia Pudelko-Malik, Dominika Drulis-Fajdasz, Mateusz Fydryszewski, Shawn Burgess, Piotr Mlynarz, Dariusz Rakus, Stanislaw Deja

**Affiliations:** Department of Biochemistry, Molecular Biology and Biotechnology, Faculty of Chemistry, Wroclaw University of Science and Technology, Wybrzeże Wyspiańskiego 27, 50-370, Wroclaw, Poland; Department of Molecular Physiology and Neurobiology, University of Wroclaw, Sienkiewicza 21, Wroclaw, 50-335, Poland; Center for Human Nutrition, University of Texas Southwestern Medical Center, Dallas, TX 75390-9046, USA; Department of Biochemistry, University of Texas Southwestern Medical Center, Dallas, TX 75390-9046, USA

## Abstract

Glucose is a predominant fuel for the brain supporting its high energy demand associated with neuronal signaling and synaptic activity. Long-term potentiation (LTP) is required for learning and memory formation by generating long lasting increase in synaptic strength and signal transmission between two neurons. While the electrophysiological bases of LTP are well established, much less is known about the metabolic demands of neurons involved in LTP. Common protocols used to examine synaptic activity rely on high glucose concentrations which are far from physiological glucose levels found in the brain. Here we used primary hippocampal neurons cultured under physiological (2.5 mM) and high (25 mM) glucose to investigate the metabolic effects of chemically induced LTP. Physiological glucose was associated with neuronal survival while high glucose promoted “PAS granule” accumulation. Changes in glucose altered extracellular lactate and pyruvate concentrations and affected key intracellular metabolic intermediates and neurotransmitter levels in neuronal cells without depleting the TCA cycle. LTP induction was comparable, but mitochondrial and neurotransmitter response to LTP was differentially affected physiological and high glucose conditions. Glycogen phosphorylase inhibition had minimal effects in physiological glucose but impaired synaptic responses and altered metabolite dynamics in high glucose. Our findings demonstrate that neuronal mitochondrial metabolism is closely linked to synaptic plasticity and highlight the importance of studying neurophysiological activity physiologically relevant glucose conditions.

## INTRODUCTION

The brain is an organ with high energy demands, consuming ∼20% of circulating glucose at basal metabolic rate, despite constituting only ∼2% of human body weight (Erbslöh et al., 1958). Consequently, glucose is a dominant oxidative fuel for brain cells. The majority of energy consumed in the brain is used for neuronal signaling, including maintenance of the resting potential and propagation of action potentials (Harris et al., 2012). Additionally, glucose is required for biosynthetic functions, as the entry of neuroactive compounds into the brain is restricted by the blood brain barrier (BBB). Therefore, glucose metabolism is essential for enabling and maintaining synaptic activity. Thus, neuronal regulation of glucose metabolism plays a critical role in memory loss, neurodegeneration diseases (Han et al., 2021), and neuropathies (Freeman et al., 2016).

Long-term potentiation (LTP) and long-term depression (LTD) are essential cellular mechanisms underlying learning and memory formation (Bear & Malenka, 1994). LTP produces a long-lasting increase, while LTD results in a long-lasting decrease in synaptic strength and signal transmission between two neurons. *In vitro*, LTP can be induced either by electrical or chemical stimulation (Abrahamsson et al., 2016), (Chen et al., 2011). The latter paradigm is based on co-application of glycine (N-methyl-D-aspartate receptor (NMDAR) co-agonist) and strychnine (glycine receptor (GlyRs) selective antagonist) (Chen et al., 2011), which leads to Ca^2+^ influx through NMDAR, activation of the Calcium/Calmodulin-dependent protein kinase II (CaMKII) cascade, and subsequent expression of immediate early genes including c-Fos (Lledo et al., 1995). While the electrophysiological bases of LTP are well established, much less is known about the metabolic demands of neurons involved in long term potentiation patterns.

Although brain cells consume substantial amounts of glucose, the role of glucose as a primary energy source for neurons in higher-order brain functions, such as the maintenance of synaptic strength through LTP, remains a topic of ongoing debate. Studies suggest that neurons utilize glucose either directly via glycolysis or rely on lactate produced from glucose by astrocytes (glial cells) as an energy source (Pellerin & Magistretti, 1994), (Li et al., 2023), (Suzuki et al., 2011). Either way, oxidative carbon originates from glucose, consistent with high glucose uptake by the brain. However, the interplay and energetic consequences of these two mechanisms are vastly different, which could explain in part how memory forming processes are regulated at the metabolic level. Furthermore, the cerebral glucose pool is partially buffered by brain glycogen, allowing the control of carbohydrate utilization independently of glycemia (Rich et al., 2019). Brain glycogen is predominantly located in astrocytes (Brown, 2004), although low levels of this multibranched polysaccharide has been detected in neurons (Oe et al., 2016), (Saez et al., 2014). In brain tissue, glycogen turnover occurs at normoglycemia, consistent with its role as an important local energy buffer for astrocytes, which is mobilized by functional activation or energy deficits (Duran et al., 2019), (Waitt et al., 2017). However, the extent to which glycogen plays a role in memory formation remains to be determined (Suzuki et al., 2011), (Gibbs et al., 2006), (Drulis-Fajdasz et al., 2023), (Pudełko-Malik et al., 2024).

In neurobiology, *in vitro* cultures of primary neurons are a valuable tool to selectively investigate neuroplasticity, neurotoxicity and neuronal metabolism (Giordano & Costa, 2011), (Gebril et al., 2016), (Brand et al., 1992), (Bouzier-Sore et al., 2003). Yet, neuronal culture conditions exhibit significant variability, particularly when it comes to glucose concentrations used in media formulations. Many *in vitro* neurophysiological studies utilize media with 25 mM or 30 mM glucose, in strong contrast to the 1.5 and 3 mM glucose found in cerebrospinal fluid (CSF) and the fact that brain tissue glucose remain only approximately 20% of that in arterial plasma (Madsen et al., 1999). Consideration of glucose concentration is important as high glucose was shown to be detrimental to neuron viability. Neurons exhibit lower viability and elevated number of apoptotic nuclei in 50 mM glucose compared to those cultured under 25 mM glucose (Zhao et al., 2018). Similarly, hippocampal neurons exposed to 50-150 mM glucose for 96 hours develop diabetic cognitive impairment and show an increase in reactive oxygen species (Liu et al., 2014). Thus, glucose levels significantly exceeding physiological concentrations may be of limited use in neuronal research.

Here, we used primary hippocampal neurons to investigate the influence of mitochondrial metabolism on synaptic plasticity and its dependence on high (25 mM) and low (2.5 mM) glucose concentration. Chemical LTP stimulation was successful under both conditions but changes in mitochondrial and neurotransmitter metabolism showed distinct patterns depending on media glucose concentration. Glycogenolysis inhibitor BAY U6751 showed minimal metabolic effects while changes in LTP were observed. Our observations indicate that under physiological brain glucose concentration, LTP occurs, but is associated with different metabolic responses compared to commonly used hyper physiological glucose media concentrations.

## Materials and Methods

### Animals

All procedures were performed in accordance with the EU Directive 2010/63/EU standards for animal research and received approval from the II Local Scientific Research Ethical Committee at Wroclaw University of Environmental and Life Science (approval number WBN.464.2.2020.IR). The study is reported following the Animal Research: Reporting of *In Vivo* Experiments (ARRIVE) guidelines.

### Isolation of primary neuronal cultures

Primary neuronal cultures were isolated from the hippocampal tissue of newborn SW.SWR/J mice originating from one litter. Each primary culture was based on 8 newborn hippocampi pooled together before plating. Briefly, hippocampi were isolated on ice in the dissection medium (DM) which consisted of pH: 7.4, 81.8 mM Na_2_SO_4_, 30 mM K_2_SO_4_, 5.8 mM MgCl_2_, 0.25 mM CaCl_2_, 1 mM HEPES, 0.2 mM NaOH, and 20 mM glucose. Following the dissection of the anatomical structures, the DM solution was removed and incubated at 37°C with 0.25% trypsin in 0.02% EDTA in DM, first in 1:2 ratio for 15 minutes, and secondly in 2:1 ratio for 10 minutes. Next, to stop trypsin activity, the explants were rinsed with warm MEM-FBS media consisting of standard MEM (bio-west, Bourg, France, L0440), 10% FBS (bio-west, Bourg, France, S181H), 1% MEM Non-Essential Amino Acid Solution (Merck KGaA, Darmstadt, Germany, M7145), 1% GlutaMAX Solution (Gibco, Grand Island, NY, USA, 35050), 100 U/L penicillin (Merck KGaA, Darmstadt, Germany, P4333), 0.1 mg/L streptomycin (Merck KGaA, Darmstadt, Germany, P4333), and 20 mM glucose.

The MEM-FBS medium was subsequently removed, and the explants were dissociated by gentle pipetting in fresh MEM-FBS. The cell suspension was centrifuged at 25°C at 500 x g, medium was removed, and cells were resuspended in 1.2 mL of MEM-FBS medium, plated in volume of 100 µL directly onto 9.6 cm^2^ well in a 6-well plate (GenoPlast Biotech S.A., Rokocin, Poland, 07-6006). All 6-well plates were coated 24 hours earlier with a coating solution (borate buffer: 80 mM H_3_BO_3_, 20 mM Na_2_B_4_O_7_; 2.5 µg/mL laminin, 0.1 mg/mL poly-L-lysine) and rinsed thoroughly. After 2 hours, the MEM-FBS medium was replaced with a neuronal medium consisting of Neurobasal A (ThermoFisher Scientific, Waltham, MA, USA, A2477501), 2% B27 Supplement (ThermoFisher Scientific, Waltham, MA, USA, 17504044), 0.5 mM glutamine (Merck KGaA, Darmstadt, Germany, G7513), 12.5 μM glutamate (Merck KGaA, Darmstadt, Germany, G1251), and 1% penicillin/streptomycin (bio-west, Bourg, France, L0022-100). Pure neuronal monoculture was obtained by addition of 2.5 µM Ara-C (Merck KGaA, Darmstadt, Germany, C6645) after 48 hours from isolation. Cells were cultured in 37 C in 80% humidity with 5% CO_2_ (Forma™ Steri-Cycle™ i160 incubator, ThermoFischer Scientific, Waltham, MA, USA). The purity of monocultures was evaluated by immunodetection of MAP2, a neuron-specific protein that stabilizes microtubules in the dendrites of postmitotic neurons (Soltani et al., 2005), and glial fibrillary acidic protein (GFAP), a marker of astrogliosis (Hol & Pekny, 2015) (**Figure S1**).

### Chemical induction of long-term potentiation (LTP) in primary neurons

Glucose in the neuronal medium was set either to 2.5 mM to mimic physiological brain glucose concentration or to 25 mM as is commonly employed in primary neuronal cultures. On day 13, cultured neurons were treated with 10 µM of glycogen phosphorylase inhibitor solution (BAY U6751) (Santa Cruz Biotechnology, Dallas, Texas, USA). The incubation time with BAY inhibitor was based on previous assessments of this compound stability (Pudelko-Malik et al., 2022). Chemical induction of LTP (molecular and cellular mechanism of learning) was performed on 14-day-old neuronal culture with or without addition of 10 µM of BAY inhibitor. Prior to the LTP induction, the cultures were transferred to the 37°C Ringer solution (pH = 7.3) consisted of: 137 mM NaCl, 5 mM KCl, 1.89 mM, CaCl_2_, 10 mM HEPES and either 2.5 mM or 25 mM glucose. The LTP was induced according to the protocol described previously (Chen et al., 2011) by addition of 1 µM strychnine (final concentration in medium), to inhibit glycinergic receptors, and 200 µM glycine solution (final concentration in medium), acting as a co-agonist of NMDAR (conditions referred to as +LTP). The controls consisted of neuronal cultures without the addition of glycine and strychnine (referred to as -LTP). The neuronal cultures were incubated for 60 min to allow for a metabolic shift induced by LTP stimulation. After this time, medium was collected, and immediately frozen at liquid nitrogen and stored at −80°C.

### Neuronal metabolite extraction

Prior to the extraction, each cell-well was rinsed twice with warm PBS containing 10 mM d-mannitol to maintain osmotic pressure. Next, plates were placed on ice and extracted with 2 mL of prechilled 80% methanol (Chemsolve, Lodz, Poland, 67561, 7732185). Cells were thoroughly scrapped, and solution was transferred to a new vial. The extraction step was performed twice to ensure maximum recovery. Next, extracts were sonicated on ice for 30 seconds twice. Samples were centrifuged for 10 min, at 500 x g in 4°C, supernatant was dried down and subjected for mass spectrometry analysis. The remaining cell pellet was used for total protein concentration measurements using BCA assay.

### Intracellular organic acids derivatizations for GC-MS measurements

Dried cellular extracts were mixed with 20 µL of an internal standard mixture (Cambridge Isotope Laboratories, Inc., Tewksbury, MA, USA), and 1 mL of ultrapure water, followed by sonication for 10 minutes. The samples were then centrifuged at 4°C for 10 minutes at maximum speed, and 750 µL of the supernatant was collected and dried under nitrogen at 45°C. Subsequently, 50 µL of a 2% solution of methoxylamine hydrochloride (MOX) (Merck KGaA, Darmstadt, Germany, 226904) in pyridine was added to the dried polar phase, vortexed, and incubated at 37°C for 90 minutes. Next, 50 µL of a silylation reagent, MTBSTFA (Merck KGaA, Darmstadt, Germany, 394882) was added, and the mixture was incubated at 60°C for 30 minutes. After incubation, the samples were centrifuged at 4°C for 10 minutes at maximum speed. Finally, 75 µL of the supernatant was transferred to GC vials containing a glass insert and analyzed.

### Extracellular organic acids derivatizations for GC-MS measurements

The sample preparation procedure for GC-MS analysis varied depending on the biological material analyzed, specifically 14-day neuronal culture medium or Ringer’s solution. For the cell culture medium, 20 µL of the medium was mixed with 20 µL of an organic acid standards mixture. The extraction was then performed using 80 µL of a solution consisting of acetonitrile and methanol in a 1:1 (v/v) ratio, and 0.5% formic acid (Lu et al., 2024). The samples were centrifuged at maximum speed for 10 minutes at 4°C. Then, supernatant was collected and evaporated under a nitrogen stream. The dried samples were then dissolved in 1 mL of pure water, and chromatographic separation was performed on a column. To elute all organic acids, the column was washed three times with water. Next, the aqueous fractions were lyophilized. The final steps included derivatization using a 2% solution of MOX and MTBSTFA, as described in the previous paragraph. For Ringer’s solution, the protocol differed in the volume of the sample and internal standards used. A 200 µL aliquot of Ringer’s solution was combined with 4 µL of an internal organic acid standards mixture (describe in GC-MS Instrumentation and Parameters).

### GC-MS Instrumentation and Parameters

Analysis of organic acids in hippocampal neuronal cell line extracts directly spiked with the water mix of label internal standards from Cambridge Isotope Laboratories (Inc., Tewksbury, MA, USA); Sodium [U-^13^C_3_] L-lactate (CLM-1579), Sodium [U-^13^C_3_] Pyruvate (CLM-2440), [^13^C_6_] Citric acid (ISOTEC Stable Isotopes, Merck, 606081), [^13^C_4_] L-Malic acid (CLM-8065), [^13^C_5_] 2-Ketoglutaric acid (Merck, 900568), [^13^C_4_] Fumaric acid (Merck, 606014), [^13^C_4_] Succinic acid (CLM-1574).

Gas chromatography was performed on an HP-5MS column (30 m × 0.25 mm, 0.25 µm; Agilent J&W) using the following temperature gradient: 120°C for 2 minutes, ramped at 6°C/min to 210°C, further ramped at 25°C/min to 250°C (held for 1 minute), and finally ramped at 25°C/min to 275°C and held for 5 minutes. A 1 µL injection volume was used for all samples, with a split ratio of 5:1.

Mass spectrometry analysis was carried out in SIM (selected ion monitoring) mode, targeting the following m/z ranges: 233.2–267.1 for lactate, 174.1–180.1 for pyruvate, 231.1–237.1 for succinate, 200.1–296.1 for fumarate, 346.2–354.2 for malate, 287.2–294.1 for oxaloacetate, 332.2–338.2 for α-ketoglutaric acid, and 571.40–576.40 for citrate. Absolute quantification was performed by comparing the peak areas of individual ions to those of internal standards based on calibration curves using MassHunter software (Agilent Technologies, Inc., USA).

### Amino acids derivatizations for LC-MS measurements

Dried cellular extracts were directly spiked with water containing a mixture of labeled amino acid internal standards (Cambridge Isotope Laboratories, Inc., Tewksbury, MA, USA): Metabolomics amino acid mix (MSK-A2-1.2), [3,3,4,4,5,5-²H₆] L-Ornithine (749443), [4,4,5,5-²H₄] L-Citrulline (DLM-6039), [¹³C₄,¹⁵N₂] L-Asparagine (641952), [Indole-²H₅] L-Tryptophan (DLM-1092), [¹³C₅] L-Glutamine (605166), [¹³C,¹⁵N₂] Urea (CNLM-234), and [¹³C₄] GABA (8666-0.05). After centrifugation at 4°C for 10 minutes at maximum speed, the supernatant was collected and dried under nitrogen. Subsequently, 100 µL of 3N HCl in n-butanol (Regis Technologies, Inc., Morton Grove, IL, USA, 7647010) was added to the sample, vortexed, incubated at 65°C for 15 minutes, and evaporated to dryness at room temperature under nitrogen. After the derivatization reaction, samples were reconstituted in 100 µL of a 50:50 (v/v) acetonitrile solution containing 0.025% formic acid and centrifuged at 4°C for 10 minutes at maximum speed. Finally, 75 µL of the supernatant was transferred to LC vials containing a glass insert and analyzed.

### LC-MS Instrumentation and Parameters

Intracellular extracts of derivatized amino acids were analyzed using an integrated triple quadrupole mass spectrometer, API 3200 LC/MS/MS (Applied Biosystems/Sciex Instruments), with electrospray ionization (ESI). Data was collected exclusively in positive ionization mode using MRM (multiple reaction monitoring). Chromatographic separation was performed on a C18 column (Xbridge, Waters, Milford, MA; 150 × 3.0 mm, 3.0 µm) in a reversed-phase system. Phase A consisted of a 50:50 acetonitrile: water mixture with 0.025% formic acid, while phase B was a 2:98 water: methanol mixture with 0.0125% acetic acid. The gradient used was as follows: 0.10 min – 0% B, 2.00 min – 20% B, 11.00 min – 60% B, 11.50 – 17.50 min – 100% B, and 18.00 – 22.00 min – 0% B. Absolute quantification was performed by comparing the peak areas of individual ions for each analyte to the peak areas of their respective internal standards based on calibration curves.

### Protein sample quantification

The total protein concentration of cell pellets was assessed using BCA protein assay kit (ThermoFisher Scientific, Waltham, MA, USA, 23225). Briefly, the cell pellet was dissolved in 1% RIPA buffer (Cell Signaling Technology, Danvers, MA, USA, 9806S) solution, vortexed and left at 4°C overnight. Next, samples were vortexed and centrifuged at 4°C at 500 x g for 10 minutes. Then, 50 µL of supernatant was combined with 1 mL of working solution (prepared as described in the BCA protein assay kit). Following 30 min incubation at 37°C, samples were left for 10 min., at room temperature. Prior to each series of measurements, the spectrophotometer was calibrated against a blank sample. Absorbance was measured at 562 nm.

### Periodic Acid – Schiff (PAS) staining in 14 days primary neuronal culture

Glycogen in primary neuronal cultures was staining of using PAS method. Briefly, cells were fixed in the solution of 96% ethanol (Chemsolve, Lodz, Poland, 1170), and methanol in ratio 3:1 (v/v). Next, fixing solution was substituted with cold 4% formalin (PFA) solution (Merck KGaA, Darmstadt, Germany, 30525-89-4) in ethanol, and incubated at room temperature for 5 minutes. 4% PFA solution was removed, and cells were thoroughly rinsed with tap water to remove any remaining alcohol. 1 mL of 1% periodic acid solution (Merck KGaA, Darmstadt, Germany, 395B), was added and cells were incubated in room temperature for 5 minutes. Plates were washed several times with distilled water, and 1 mL Schiff base reagent (Merck KGaA, Darmstadt, Germany, 395B) was added to the well for 15 minutes. Finally, the coverslips were washed several times with tap water and subjected to the immunofluorescence staining.

### Glucose measurement

The glucose level was measured based on Ringer solution using Glucose (HK) Assay Kit (Merck KGaA, Darmstadt, Germany; G3293). The absorbance was read at 340 nm using a Synergy H1 hybrid plate reader (Agilent BioTek, USA).

### Immunofluorescence after glycogen staining

The immunofluorescence staining was performed on the neuronal cells after PAS procedure. To avoid unspecific binding of the primary antibody, cells were incubated with 1% bovine serum albumin (BSA, Merck KGaA, Darmstadt, Germany, 9048-46-8) in room temperature for 1h. After intensive washing in PBS, cells were incubated at 37°C for 30 min, with solution containing 0.001% TritonX-100, (Merck KGaA, Darmstadt, Germany9002-93-1), 0.1% BSA, mouse anti-GFAP antibody (1:200, Merck KGaA, Darmstadt, Germany, SAB5201104), and rabbit anti-MAP2 antibody (1:200, ThermoFisher Scientific, Waltham, MA, USA, PA5-17646). After incubation period, cells were washed twice with PBS, and the fluorophore-labeled secondary antibodies were added: goat anti-mouse-AlexaFluor488 (1:2000, ThermoFisher Scientific, Waltham, MA, USA, A-11029), and goat anti-rabbit-AlexaFluor568 (1:2000, ThermoFisher Scientific, Waltham, MA, USA, A-11061). After 2.5 hours of incubation at room temperature cells were rinsed once with PBS and nuclei were visualized using DAPI (ThermoFisher Scientific, Waltham, MA, USA, A-R37606). To ensure the best visualization of glycogen in neuronal cells, the microscope slides were analyzed within 24 hours.

### Immunofluorescent studies (c-Fos, CaMKII)

For immunofluorescent staining, primary neuronal cell cultures were fixed in an alcoholic fixative (1:3 methanol: ethanol, 95%) solution for 20 minutes at 4°C and subsequently stored at −20°C. Prior to immunolabeling, the fixed cells were rehydrated in distilled water (30 minutes, room temperature [RT]) followed by phosphate-buffered saline (PBS, pH 7.4; 30 minutes, RT), and incubated with 1% bovine serum albumin in PBS for 1 hour at RT. The cells were then incubated overnight at 4°C with primary antibodies diluted in PBS containing 0.1% BSA and 0.001% Triton X-100. The primary antibodies used were anti-MAP2 (1:500; ThermoFisher Scientific, Waltham, MA, USA, PA5-17646), anti-GFAP (1:500; ThermoFisher Scientific, Waltham, MA, USA, G9269), anti-c-Fos (1:500; ThermoFisher Scientific, Waltham, MA, USA, MA1-21190), anti-CaMKII (1:500; Abcam, 1082903-4). Following primary antibody incubation, cells were incubated for 2.5 hours at RT with species-specific secondary antibodies diluted in PBS with 0.1% BSA. The secondary antibodies used included goat anti-mouse Alexa Fluor 633 (1:2000; ThermoFisher Scientific, Waltham, MA, USA, A21046), goat anti-rabbit Alexa Fluor 488 (1:2000; ThermoFisher Scientific, Waltham, MA, USA, A11034), goat anti-rabbit Alexa Fluor 633 (1:2000; ThermoFisher Scientific, Waltham, MA, USA, A21071), and goat anti-mouse Alexa Fluor 488 (1:2000; ThermoFisher Scientific, Waltham, MA, USA, A11001), and goat anti-chicken Alexa Fluor 568 (1:2000; Abcam, ab175477). Finally, glass slides were mounted with Fluoroshield containing DAPI (cat. no. F6057, Sigma-Aldrich). Between each step of the immunostaining procedure, cells were washed three times with PBS (10 minutes per wash, RT).

### IF image analysis

PAS based glycogen visualization was carried out using Olympus BH-2 Polarizing Trinocular Microscope (DPlan 20x/0.40 NA). Immunofluorescence image analysis of MAP2, c-Fos and CaMKII (pT286CaMKII) protein were conducted using the Confocal Laser Scanning Microscope FV3000 (Olympus) with 60× objective, at 1024 × 1024 picture resolution, with identical acquisition parameters set individually for each analyzed protein. ImageJ software was used for the quantification of the mean fluorescence intensity (Schneider et al., 2012). The mean fluorescence intensity was measured from regions of interest (ROIs) and normalized to cell area.

### Statistical Analysis

Univariate analysis was performed GraphPad Prism (version 10). Data are expressed as the mean ± SEM, with the sample size (n) for each group displayed as individual points or indicated in the corresponding figure legends. Depending on data distribution Student’s t-test or Mann-Whitney-Wilcoxon test was applied. The data distribution of the results was evaluated using the Shapiro-Wilk test. All statistical tests were conducted at a significance level of ^∗^p < 0.05, ^∗∗^p < 0.01, ^∗∗∗^p < 0.001.

## Results

### Primary neuronal cells are viable when cultured in low glucose media

Glucose concentrations in the range of ∼15 – 25 mM are commonly used for *in vitro* neuronal cultures, despite being 10 times higher than the physiological glucose concentration found in the brain (Madsen et al., 1999). For that reason, alternative protocols utilizing glucose concentrations closer to physiological levels have been proposed (Kleman et al., 2008), (Kamal et al., 1999), (Bardy et al., 2015). Thus, we aimed to test if the glucose concentration in media impact neuronal cell count over 14 days of culture (**Figure 1A**). The same volume of cell suspensions isolated from an equal number of hippocampi were plated at the beginning of each experiment. After 14 days cell nuclei were stained with DAPI to visualize and quantify the cells. A greater density of DAPI-stained nuclei was observed in 2.5 mM compared to 25 mM glucose suggestive of a higher cell count (**Figure 1B**). Since primary neurons are unable to divide *in vitro*, the changes in their number correspond to their survival. Quantitative analysis confirmed a significant decrease in the number of DAPI-stained nuclei in cultures grown under 25 mM relative to 2.5 mM glucose (**Figure 1B**). These data suggest that low media glucose may either promote neuronal survival, or high media glucose may exert a cytotoxic effect during extended culture period.

**Figure 1.**
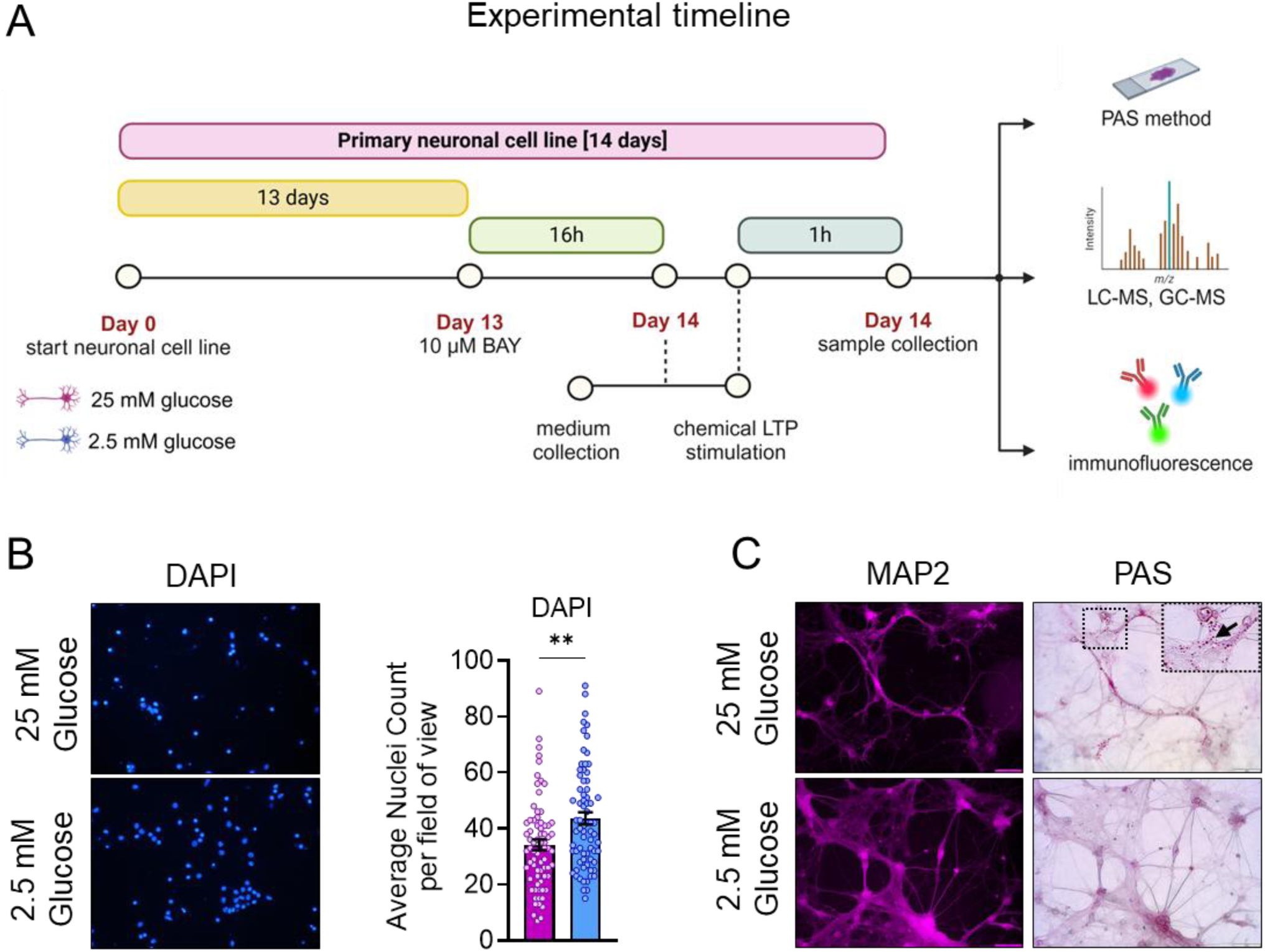
Impact of glucose concentration on viability and presence of PAS granules in extended neuronal cells. (**A**) Experimental design conducted on 14-day-old neuronal cultures incubated in either 25 mM or 2.5 mM culture medium. Chemical LTP stimulation was applied for one hour to the cultures. Samples were analyzed by LC-MS to quantify intracellular amino acids, while GC-MS was used to determine intracellular and extracellular cellular acid intermediates. Samples were also assessed by immunofluorescence for the detection of MAP2, GFAP, c-Fos, and p-CaMKII proteins. (**B**) Neuronal cell count assessed by DAPI staining in relation to glucose concentration in the culture medium (n = 72, 25 mM glucose, n = 72, 2.5 mM glucose). (**C**) Detection and localization PAS granules within neuronal cells in response to different glucose concentrations in the culture medium (n = 64, 25 mM glucose, n = 61, 2.5 mM glucose). Data are presented as mean ± SEM with the (n) of each group shown as individual points. Two group comparisons were evaluated using an unpaired Student’s t test or Mann-Whitney’s test depending on the data distribution. Significance is given as: *p < 0.05, **p < 0.01, ***p < 0.001.

### Presence of PAS granules in primary neuronal cells is dependent on the media glucose concentration

Cellular glycogen is influenced by extracellular glucose concentrations. Although both neurons and astrocytes are capable of glycogen storage, the majority of brain glycogen is located in astrocytes (Brown, 2004). Indeed, the mouse brain stains positively with Periodic Acid-Schiff (PAS) which is not specific to glycogen but can be associated with it (Akiyama et al., 1986). Additionally, “PAS granules” can be detected in hippocampus and other regions of the brain, but glycogen is not their main component (Manich et al., 2016). PAS granules have been identified in brain tissues and associated with neurodegenerative diseases, but their cellular distribution and origin are not clear.

By isolating primary neurons, we were able to conduct PAS staining in cell specific manner. Neuronal purity of culture was evaluated using MAP2 staining (a neuron specific protein marker), which colocalized with PAS to confirm specificity to neuronal cell bodies (**Figure 1C**). Astrocytes were sparsely detected (below 10%), but only images containing neurons without astrocytes were evaluated. Qualitative evaluation of fluorescence microscopy images revealed a pronounced accumulation of “PAS granules” in neuronal cells cultured in 25 mM glucose condition, as indicated by the intense clustered PAS staining (**Figure 1C, black arrow**). In contrast, neuronal cells maintained in 2.5 mM glucose conditions displayed minimal or no observable “PAS granules” despite considerable PAS staining dispersed within cells. This data suggests that high media glucose concentration promotes “PAS granule” accumulation in primary neurons, while low glucose conditions prevent their development. Due to the non-specific nature of PAS the extent to which neuronal glycogen or glycoproteins contributes to these observations remains to be determined.

### Low media glucose depletes intracellular neurotransmitters and their metabolites in primary neurons

Given the differential effects of 25 mM and 2.5 mM glucose concentration on cell number and polysaccharides distribution (PAS staining), we hypothesized that intracellular neuronal metabolite levels would also be dependent on glucose availability. Since neurons sustain a high rate of oxidative metabolism to support synaptic activity (Magistretti & Allaman, 2015), we focused on pyruvate, TCA cycle intermediates and neurotransmitters corresponding to major neuronal metabolic function. Quantitative targeted metabolomics results were normalized to total protein content to account for differences in cell count observed between glucose conditions.

We detected decreased levels of intracellular pyruvate and its metabolites, lactate and alanine, consistent with limited glycolysis under 2.5 mM compared to the 25 mM glucose media (**Figure 2A**). Notably, there were no changes in lactate/pyruvate (L/P) ratio, indicating that depletion of pyruvate related metabolites occurred without changes in cytosolic redox state (**Figure 2A**). Consistent with cellular data, the extracellular lactate and pyruvate concentrations were also significantly lower, indicating less glucose fermentation (**Figure 2B**).

**Figure 2.**
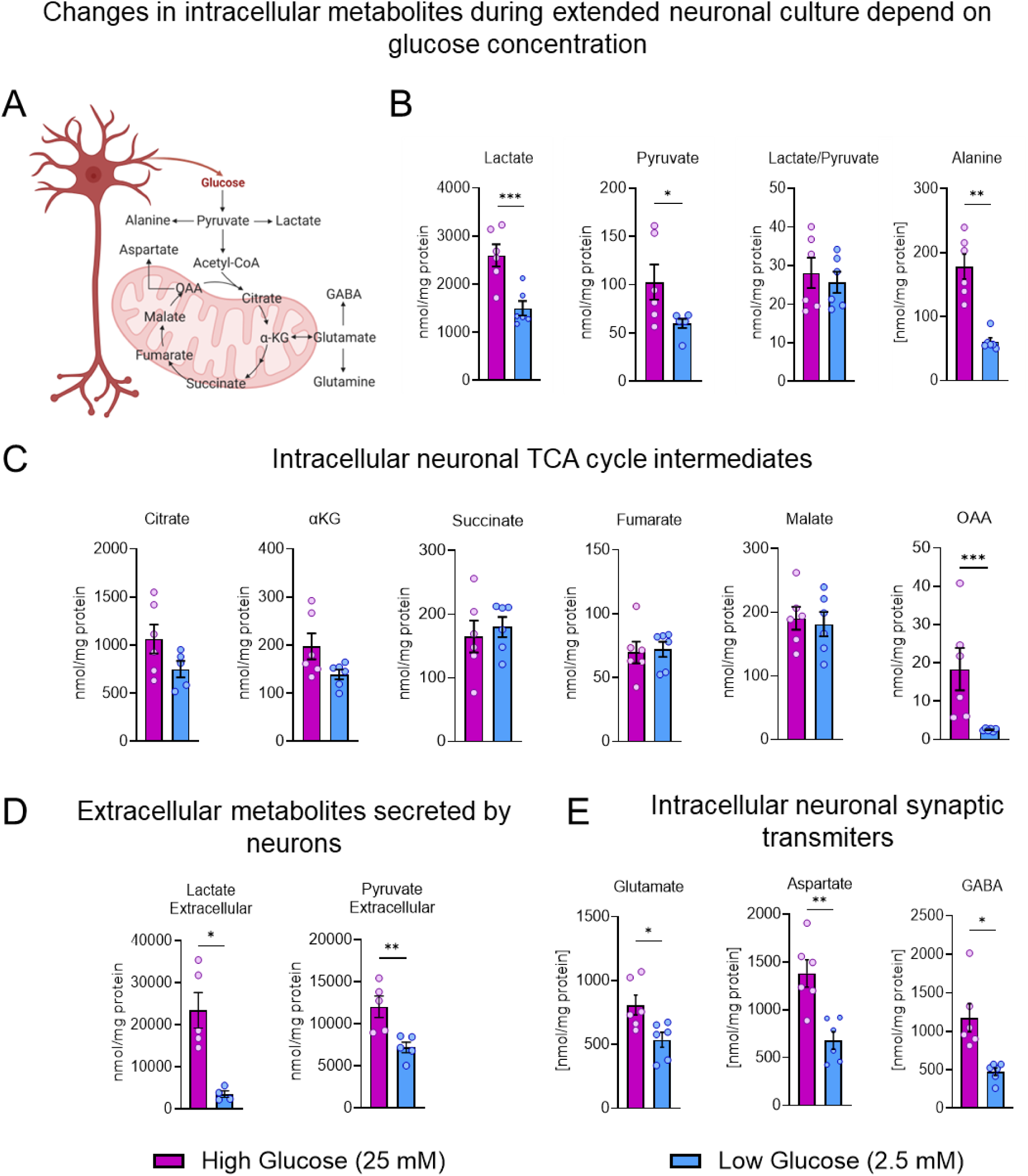
Influence of different glucose concentrations on mitochondrial metabolism and neurotransmitters in primary neuronal cells. (**A**) Schematic illustration of mitochondrial metabolism and its correlation with neurotransmitters. (**B**) Concentration of intracellular alanine and organic acids in 14-day-old primary neuronal cells. (**C**) Intracellular TCA cycle intermediate concentrations depending on different glucose concentrations. (**D**) Changes in lactate and pyruvate extracellular concentrations in extended neuronal medium. (**E**) Intracellular synaptic transmitter concentrations in response to 25 mM and 2.5 mM glucose. All intracellular metabolite measurements were based on n = 6 biological replicates, and n = 5 for medium.

Despite large changes in pyruvate, TCA cycle intermediates remained surprisingly resistant to changes in media glucose concentration. While citrate and αkg trended lower, all four carbon TCA cycle intermediates with exception of OAA (**Figure 2C**) were not different. Massive depletion of OAA correlated with a substantial decrease in aspartate suggesting that their levels are regulated independently of succinate, fumarate and malate in primary neurons. Furthermore, neurotransmitters such as glutamate, aspartate, and GABA were all significantly lower in neurons cultured in 2.5 mM compared to 25 mM glucose (**Figure 2D**). These results suggest that low glucose availability significantly alters key metabolic intermediates and neurotransmitter levels in neuronal cells without necessarily depleting the TCA cycle.

### Low glucose concentration does not impair LTP induction in 14-day primary neuronal cultures

Based on observed depletion of intracellular neurotransmitters in low-glucose media, we hypothesized that 2.5 mM glucose media would be associated with less potent LTP induction in hippocampal neurons. Instead, we detected a surprisingly strong c-Fos immunofluorescence signal following glycine-induced LTP in 2.5 mM glucose (**Figure 3B**). c-Fos is an immediate-early gene (IEG) involved in modulating neuronal plasticity and memory formation and therefore is a common marker of successful LTP stimulation (Lara Aparicio et al., 2022). The degree of c-Fos expression following LTP induction was comparable between both glucose conditions (**Figure 3A,B**). To further confirm that stimulation resulted in LTP and not LTD, we analyzed the phosphorylation of CaMKII, a critical mediator of early-phase LTP and synaptic strengthening. A significant increase in p-CaMKII fluorescence in 2.5 mM glucose compared to the non-stimulated control was observed (**Figure S2**). Taken together, these data suggest that LTP can occur *in vitro* at physiological brain glucose concentrations.

**Figure 3.**
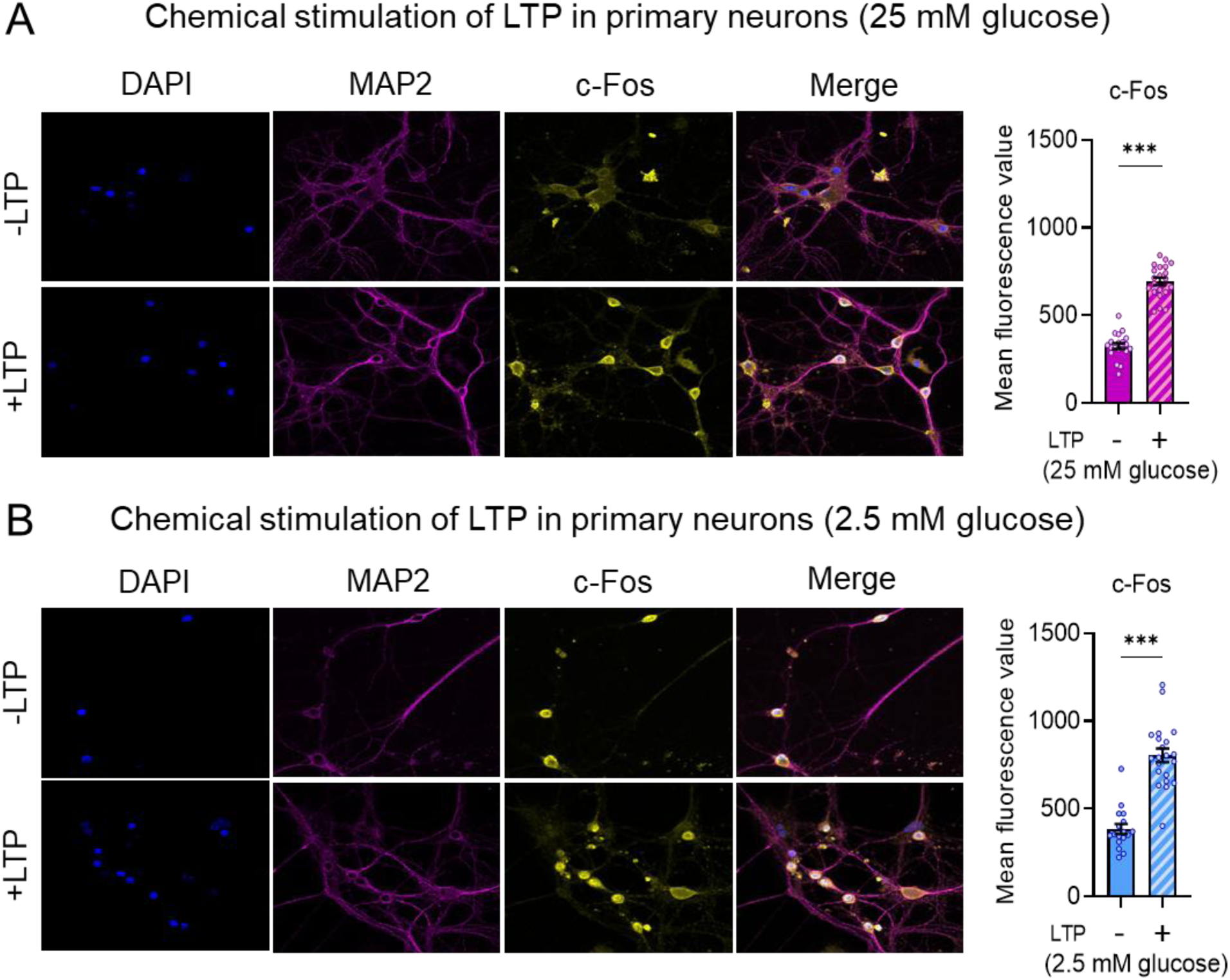
Expression of c-Fos protein in the hippocampal primary neurons is not affected by low glucose concentrations in the medium during LTP stimulation. (**A-B**) Representative confocal images of c-Fos immunofluorescence (yellow) distribution in neuronal cells 60 minutes after LTP induction using 1 µM strychnine and 200 µM glycine in Ringer solution in (**A**) 25 mM glucose and (**B**) 2.5 mM glucose. Localization of neuronal soma, dendrites and nuclei was imaged using antibodies against MAP2 (magenta), and DAPI (blue). Quantification of c-Fos immunofluorescence was calculated based mean fluorescence value and normalized to values obtained for all cel (n = 20 (25mM), n = 21 (25mM+LTP), n = 17 (2.5mM), n = 21 (2.5mM+LTP).

### Changes in neuronal metabolism in response to chemical induction of LTP depend on glucose availability

Since LTP induction was observed despite differences in glucose availability, we next investigated whether neuronal metabolism was also comparable. We quantified TCA cycle metabolites in neurons without stimulation (baseline) and following chemical induction with glycine and strychnine (stimulation). Metabolic response to LTP was expressed as a ratio 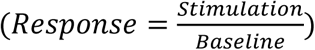 and was differentially affected by 25 mM and 2.5 mM glucose (**Figure 4A**). In 2.5 mM glucose, the intracellular TCA cycle intermediates accumulated, while in 25 mM glucose they were depleted following LTP stimulation. Notably, similar responses were observed for intracellular synaptic neurotransmitters (**Figure 4A**), highlighting their relationship with mitochondrial metabolites. During 60 minutes of LTP stimulation, lactate secretion from neurons decreased in 25 mM and increased in 2.5 mM glucose, consistent with changes in intracellular metabolites (**Figure 4A**). However, when we quantified residual glucose in the media, the response was either unchanged (Response ∼1) or slightly increased (Response ∼1.1) by 25 mM and 2.5 mM glucose, respectively. Neurons cannot produce glucose, thus the increased glucose concentration after LTP stimulation compared to baseline indicates less glucose uptake in 2.5 mM glucose. On the other hand, no observable effect in 25 mM glucose, may be due to the analytical limitation in detecting small differences in glucose uptake when media glucose concentration is high (25 mM).

**Figure 4.**
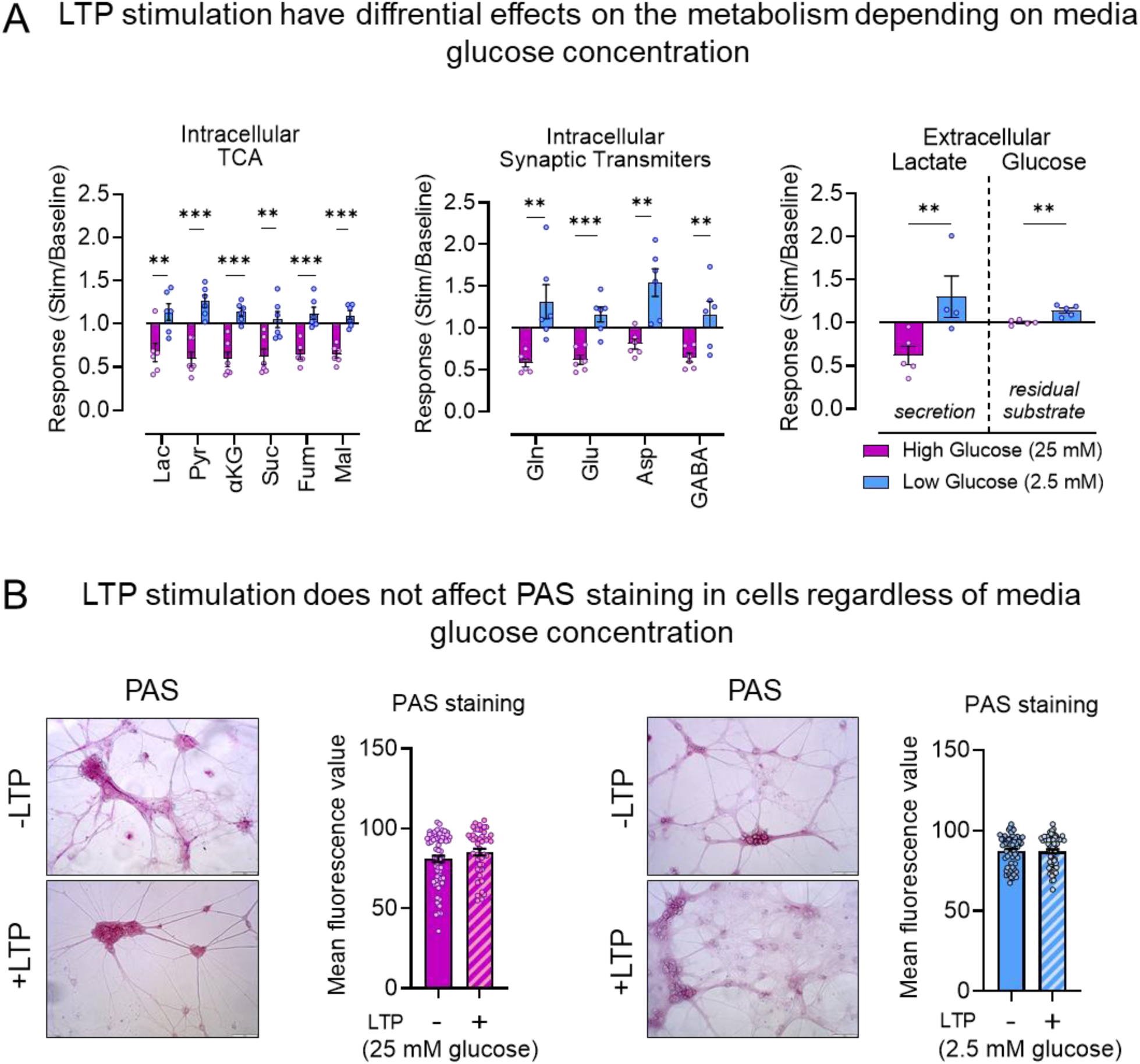
Induction of LTP in low glucose increase response of TCA intermediates and neurotransmitters. (**A**) Intracellular and extracellular metabolic response to long term potential stimulation in low (2.5 mM), and high (25 mM) glucose (n = 6 (25 mM), n = 6 (25 mM + LTP), n = 6 (2.5 mM), n = 6 (2.5 mM + LTP). (**B**) Quantification of PAS staining within neuronal cells after 60 minutes of LTP stimulation in the Ringer medium with 25 mM or 2.5 mM glucose. (n = 64, 25 mM glucose; n = 61, 2.5 mM glucose). Due to the intensity of staining (granules in high glucose) and their absence in low glucose, it was necessary to apply different threshold values to cover the entire cell area. The threshold used for quantification in high glucose was approximately 70, whereas in low glucose, it was around 95. Therefore, quantification values cannot be directly compared between glucose concentrations.

The observed changes in intracellular and extracellular metabolism during LTP stimulation highlight a potential disconnect between intracellular metabolite response and glucose uptake, theoretically implicating the involvement of glycogen pool. To investigate this possibility, we aimed to examine PAS staining in neurons during chemical long-term potentiation (LTP) stimulation. Our results indicated no differences in PAS staining between pre-stimulation and post-stimulation conditions, regardless of the glucose concentration (**Figure 4B**) indicating that LTP induction does not lead to measurable changes in PAS positive carbohydrate features of neurons. Due to the non-specific nature of PAS, we cannot conclude that there were no changes in glycogen, but the overall effect of LTP induction on PAS staining was minimal.

### Inhibition of glycogenolysis affects LTP induction under high glucose conditions

Although there were no detectable changes in PAS staining during LTP, it remains possible that glycogen metabolism is required for LTP. As mentioned above, the PAS specificity is not limited to glycogen, and the inhibition of glycogen phosphorylase was shown to modulate synaptic plasticity in aged mice (Drulis-Fajdasz et al., 2018), (Drulis-Fajdasz et al., 2015). To test the potential role of neuronal glycogen in LTP stimulation we applied the glycogen phosphorylase inhibitor BAY in 25 mM and 2.5 mM glucose incubation conditions a day before and during LTP stimulation (**Figure 1A**). In 2.5 mM glucose the chemical induction of LTP in the presence of 10 µM BAY resulted in substantially increased c-Fos levels, similar to the control condition without BAY (**Figure 5A**). Minimal effects on glycogenolysis inhibition was expected, since the amount of glycogen in neurons is especially low when incubated in 2.5 mM glucose. Surprisingly, 10 µM BAY increased c-Fos substantially even at baseline when neurons were incubated in 25 mM glucose. Furthermore, we did not observe the induction of c-Fos following the chemical induction of LTP in 25 mM glucose when BAY was present (**Figure 4A**). These findings suggest that BAY alone can affect neuronal genes expression in a glucose dependent manner and suggest that mechanisms of synaptic plasticity could be modulated by inhibition of glycogen phosphorylase (**Figure 4B**).

**Figure 5.**
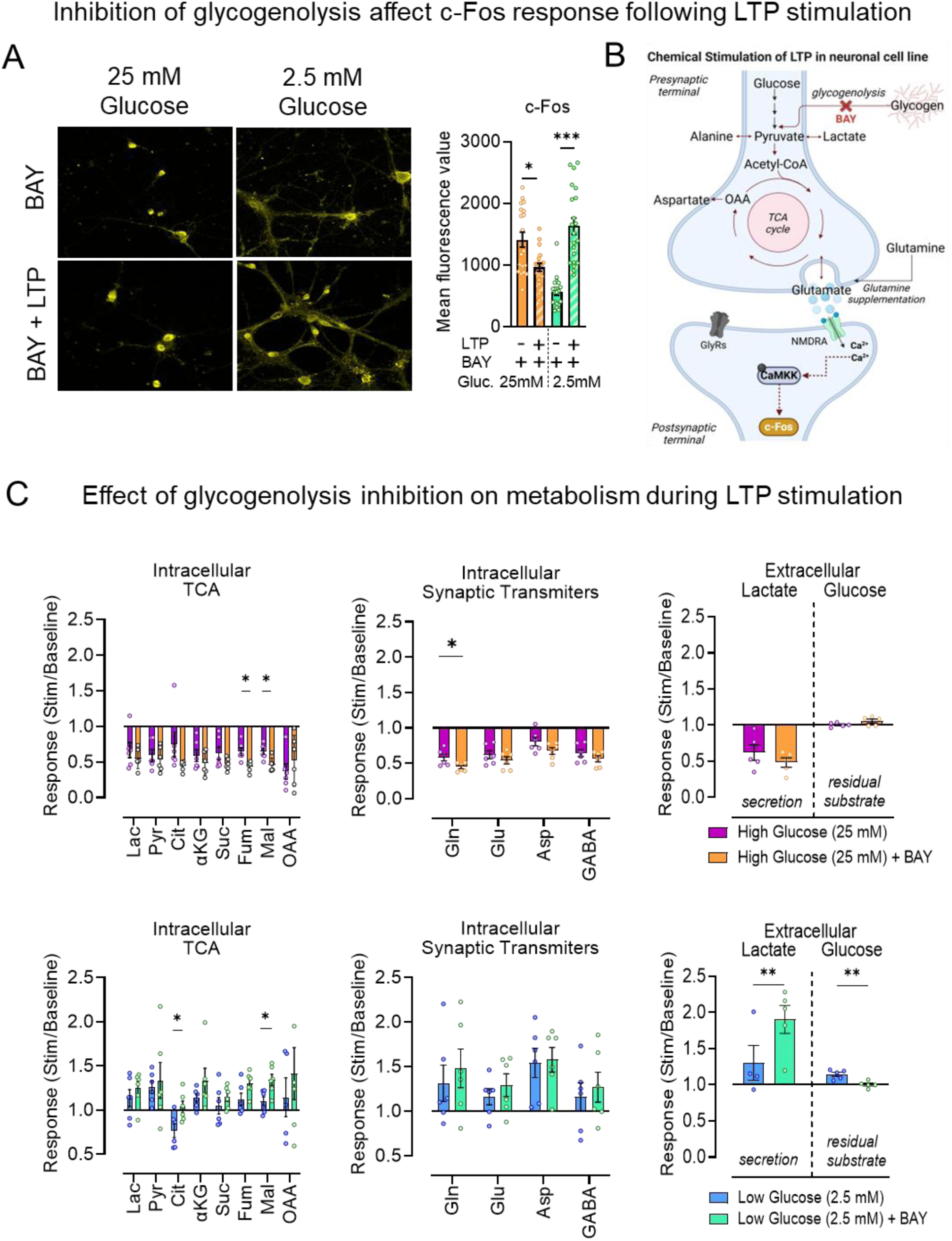
Inhibition of glycogenolysis in high glucose impairs synaptic transmission and neuronal metabolism. **(A)** Representative confocal images of c-Fos immunofluorescence (yellow) distribution in neuronal cells 60 minutes after of glycine-induced LTP, with the addition of 10 µM BAY, an inhibitor of glycogen phosphorylase, in Ringer’s solution at two different glucose concentrations (25 mM and 2.5 mM). Quantification of c-Fos immunofluorescence was performed based on the mean fluorescence intensity and normalized to the values obtained for all cells (n = 19 for 25 mM + BAY, n = 20 for 25 mM + BAY + LTP, n = 20 for 2.5 mM + BAY, n = 22 for 2.5 mM + BAY + LTP). (**B**) Scheme illustrating the mechanism of neuronal metabolism with inhibition of glycogenolysis during LTP stimulation. (**C**) Intracellular and extracellular metabolic response to long term potential stimulation under glycogen breakdown blockade (BAY) in low (2.5 mM), and high (25 mM) glucose (n = 6 for all intracellular metabolite measurements; n = 5 for extracellular lactate and glucose).

Next, we investigated the effect of BAY mediated inhibition of glycogenolysis during LTP stimulation on TCA cycle intermediates and neurotransmitters. Importantly, BAY treatment prior and during LTP stimulation (**Figure 5C**) did not change the directionality of responses to LTP for most metabolites investigated (**Figure 4A**). In 2.5 mM glucose media there was a statistically significant increase in the response of citrate and fumarate to LTP stimulation but no differences in response of neurotransmitters following BAY application (**Figure 5C**). In 25 mM glucose, BAY caused a statistically significant decrease in the response of glutamine, fumarate and malate levels. In 25 mM glucose, BAY did not change glucose consumption or lactate secretion during LTP stimulation. Conversely, in 2.5 mM glucose, BAY significantly increased lactate secretion and prevented the glucose sparing effect during LTP stimulation (**Figure 4C**). Taken together this data suggests that inhibition of glycogenolysis may affect neuronal metabolism during LTP stimulation, but the degree of this effect is dependent on extracellular glucose concentration.

## Discussion

Blood glucose concentration is linearly correlated with glucose levels found in the brain and its transport through the BBB follows Michaelis-Menten kinetics as evidenced by in vivo ^1^H and ^13^C MRI measurements in humans (Gruetter et al., 1998). Brain glucose concentration can be estimated to be ∼2.5 mM, based on MRI estimates of 1-8 µmol/g (Gruetter et al., 1998) and the brain tissue density ∼1.041 g/mL (Weaver et al., 2001). However, standard electrophysiology protocols differ vastly in the concentration of glucose in the artificial cerebrospinal fluid (ACSF), which range from 0.5 to 25 mM (van den Top et al., 2017), (Kay & Krupa, 2001), (Abrahamsson et al., 2016). In primary neuronal cultures, the 25 mM glucose is commonly used (Meng et al., 2014), with even higher media glucose concentrations (e.g., 50, 75, or 150 mM) in some cases (Liu et al., 2014). These non-physiological concentrations cannot mimic normal neuronal function, as such high glucose concentrations (e.g. ∼12 mM) would only be achieved with blood glucose concentrations of approximately 60 mM (∼1000 mg/dL), an extreme hyperglycemia. In human, this would cause a loss of consciousness and diabetic coma, as diabetic hyperosmolar syndrome occurs when blood glucose level exceeds 33.3 mM (600 mg/dL) (Kitabachi et al., 2014). Therefore, more attention needs to be directed towards concentrations of nutrients in neuronal experimental media that mimic normal physiology, or within the limits of pathophysiology (e.g., higher glucose concentrations may be relevant e.g. when designing experimental paradigms focused on diabetes and related neuropathies). Nevertheless, we found that different glucose concentrations in the media of primary neurons are sufficient to not only change the mitochondrial metabolite concentrations in these cells but also affect the degree to which they respond to chemical LTP stimulation. The exact nature of this phenomena requires more investigation in the future.

Electrophysiological studies investigating hippocampal slices indicate that glucose concentration in ACSF influences the baseline excitatory post-synaptic potentials (Kamal et al., 1999) but has no effect on neurotransmitter release (Kamal et al., 1999). More specifically, a minimal amount of glucose is essential for LTP to occur, as tetanization of slices perfused with 0 mM glucose results in no changes in the slopes of the fEPSPs (Kamal et al., 1999). Conversely, LTP induction is enhanced with increasing glucose in media, and appears to a reach maximum at 10 mM, as 30 mM glucose does not it improve further (Kamal et al., 1999). Therefore, we conclude that in the physiological range of brain glucose (0.5 – 3.0 mM), the LTP indeed can take place but is also associated with the maximal variability in terms of its effect size. Thus, it could be that glucose, and its downstream metabolite pyruvate, are important regulators of the degree to which LTP takes place. Our data is consistent with these observations as we were able to obtain substantial c-Fos induction in 2.5 mM, but the induction was even stronger in 25 mM glucose. Pyruvate, lactate secretion and mitochondrial metabolism were differentially affected during LTP induction in 2.5 mM and 25 mM glucose, highlighting diverse partitioning of pyruvate depending on glucose availability.

There are at least three roles that glucose can serve during the LTP event: 1) supporting large amounts of pyruvate for oxidative phosphorylation and generation of ATP, as LTP has been postulated to require large amounts of energy, 2) providing a carbon backbone for neurotransmitters and protein synthesis, and 3) contributing to glycogen stores that control intracellular metabolites availability needed to facilitate neuronal plasticity.

The baseline energetic requirements do not appear to be a factor as, the ATP neuronal content remains unchanged across different glucose concentrations (0.2, 5 and 10 mM) (Fleck et al., 1993). However, neuronal oxidative metabolism following stimulation and more importantly, the transition to the consolidation (maintenance) phase of LTP may be dependent on ATP (Wieraszko & Ehrlich, 1994). We did not measure ATP levels, however in low glucose media there was an accumulation of TCA cycle intermediates and increased lactate release with a simultaneous sparing of glucose in the media. This data does not necessarily support enhanced glycolysis nor increased oxidative TCA cycle turnover but is rather consistent with induction of anaplerosis or inhibition of cataplerosis in the TCA cycle and thus capturing carbon in the TCA cycle. It is possible that under such conditions, less ATP can be produced, and LTP stimulation is not as strong as observed in high glucose conditions. Conversely, in high glucose media following LTP induction there was a decrease of lactate secretion with no change to glucose uptake form the media, and TCA cycle intermediates were diminished compared to baseline prior stimulation. This data is consistent with changes in pyruvate partitioning away from LDH based lactate release in favor of oxidative pyruvate metabolism via PDH rather than anaplerotic pyruvate carboxylation. While it is difficult to infer metabolic fluxes from metabolite levels, lower TCA cycle intermediates are not preventive of high TCA cycle turnover and generation of ATP.

Changes in pyruvate metabolism were linked to alterations in potassium-evoked neurotransmitter release from neurons (Fleck et al., 1993). In the presence of 10 mM glucose, potassium stimulation enhances the release of glutamate and aspartate. However, under reduced glucose availability (0.2 mM), potassium decreases glutamate and increases aspartate release respectively (Fleck et al., 1993). These findings are consistent with our observations of increased aspartate levels in 2.5 mM glucose condition following stimulation of LTP, which would be consistent with enhanced glutamate catabolism in the TCA cycle. This metabolic shift likely prevents net glutamate release in favor of increased aspartate production.

Although brain glycogen has traditionally been considered predominantly localized in astrocytes (Brown, 2004), there is evidence for its presence within hippocampal and cortical neurons (Oe et al., 2016). Accumulation of neuronal glycogen can be neurotoxic as it has been shown to induce apoptosis and autophagy (Duran et al., 2012) or, as in Lafora disease, glycogen-derived polyglucosans can lead to epilepsy and neuronal death (Robitaille et al., 1980). Nevertheless, these reports confirm the existence of glycogen synthesis machinery in neurons but also emphasize the need for its tight control in order to prevent neurotoxicity (Duran et al., 2012). In healthy brain neuronal glycogen metabolism may be of physiological relevance during postnatal brain development (Borke & Nau, 1984) and protection from hypoxia (Saez et al., 2014). Neuronal glycogen is ten times lower than astrocytes, and thus its detection can be best accomplished using glycogen specific antibodies (Baba, 1993). Here we used periodic acid Schiff (PAS) staining which is less sensitive and less specific methodology. Nevertheless, we detected strong PAS staining attributable, to in part, to glycogen stores and glycoproteins. It is important to note that our model consisted of 14-day hippocampal primary cells, which, are considered a fully mature neurons (Kepiro et al., 2018). Mature neuronal cells are thought to accumulate more polyglucosans compared to young neurons which raises an important question regarding the potential role of glycogen metabolism in higher brain functions, such as memory formation (Gibbs et al., 2006), (Suzuki et al., 2011), (Duran et al., 2013). Nevertheless, we did not observe changes in PAS staining before and after LTP stimulation.

Finaly, the observation that neuronal metabolism can be affected by glucose concentration, may be of importance with respect to diabetes and related pathologies. Diabetes is associated with neuropathies, which leads to numerous complications. Chronic high blood glucose concentration likely induces changes in glycolysis and downstream mitochondrial pathways leading to changes in neurotransmitter biosynthesis and their proportions. It is unclear how chronically elevated glucose would affect peripheral neurons compared to those located in the brain, specifically hippocampus, as it has been recently shown that cerebrospinal fluid flow extends to peripheral nerves (Ligocki et al., 2024). Nevertheless, arteriole penetrates into endoneurium potentially supplying glucose to peripheral nerves. Indeed, changes in vasculature are correlated with development of neuropathy, implicating that blood based metabolic supply may still be of importance (Yagihashi et al., 2011). Glucose metabolism in neurons could be a component of the pathological neuronal responses to chronically elevated glucose levels.

## Author contributions

Conceptualization, D.D.-F., D.R., N.P.-M., S.D., and P.M.; methodology, N.P.-M., S.D., D.D.-F., and M.F; investigation, N.P.-M. and S.D.; resources, S.D., S.B., D.R., and P.M.; writing – original draft, N.P.-M. and S.D.; writing – review & editing, D.D.-F., D.R., M.F, N.P.-M., S.D., S.B. and P.M.; funding acquisition, S.D., P.M., DR.; supervision, S.D., S.B., D.R., and P.M. D.D.-F. All authors reviewed the manuscript.

## Acknowledgments

This research has been made possible by the Kosciuszko Foundation the American Centre of Polish Culture. Financial support was provided by the Kosciuszko Foundation (N.P.-M.), by the Polish National Science Centre: UMO-2020/37/B/NZ4/00808 (D.R.), and by National Institutes of Health (NIH) K01DK133630 (to S.D.). The authors acknowledge the UT Southwestern NORC Metabolic Phenotyping and Quantitative Metabolism Cores (NIH P30DK127984).

