## Supplementary Files for "Glucose-dependent metabolism of hippocampal primary neurons in response to chemically induced long-term potentiation"

### Supplementary Figure captions

**Figure S1.** Representative confocal microscopy image of neuronal culture. Localization of neuronal soma, dendrites and nuclei was revealed with antibodies against MAP2 (magenta), for astrocyte antibodies against GFAP (green) were used, and DAPI (blue), respectively (merge channel).

**Figure S2.** Exemplary confocal images of p-CaMKII immunofluorescence (yellow) distribution in neuronal cells after glycine LTP induction in Ringer solution in 2.5 mM glucose. Localization of neuronal soma, dendrites and nuclei was revealed with antibodies against MAP2 (magenta), and DAPI (blue), respectively (merge channel). Quantification of c-Fos immunofluorescence was calculated based mean fluorescence value and normalized to values obtained for all cell (n=20 (25mM), n=21 (25mM+LTP), n=17 (2.5mM), n=21 (2.5mM+LTP).

**Figure S3.** Quantification of PAS staining within neuronal cells 60 minutes after LTP stimulation with the addition of 10 µM BAY inhibitor, in the Ringer medium with 25 mM or 2.5 mM glucose. (n = 64, 25 mM glucose; n = 61, 2.5 mM glucose).

### Supplementary Figure S1

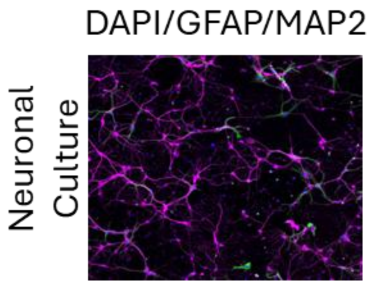


### Supplementary Figure S2

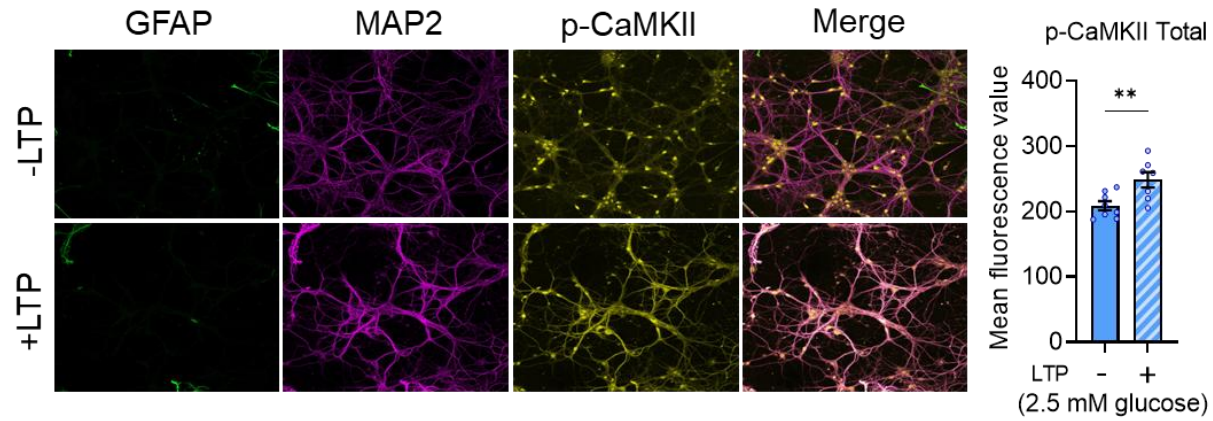


### Supplementary Figure S3

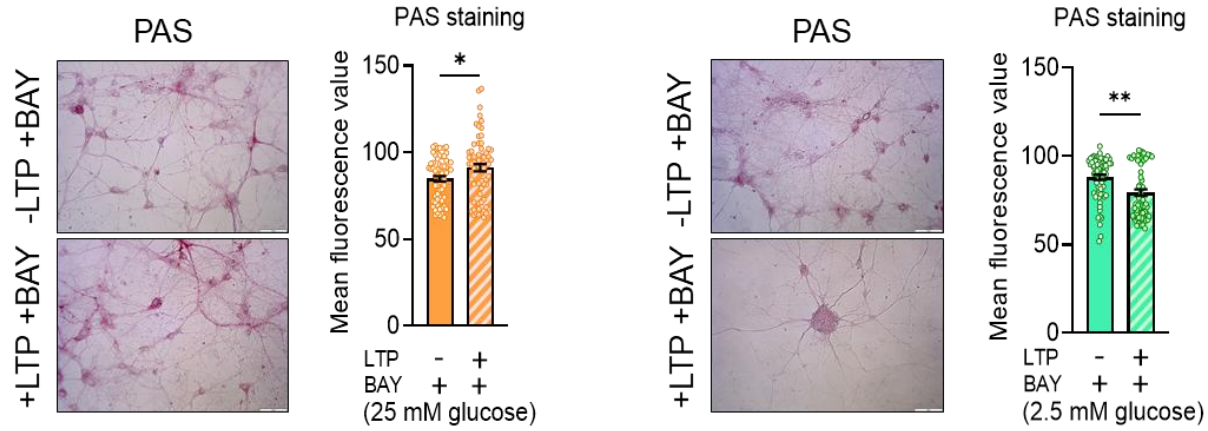
